# The responsiveness of gait and balance outcomes to disease progression in Friedreich ataxia

**DOI:** 10.1101/2021.03.18.434657

**Authors:** Sarah C Milne, Seok Hun Kim, Anna Murphy, Jane Larkindale, Jennifer Farmer, Ritchie Malapira, Mary Danoudis, Jessica Shaw, Tyagi Ramakrishnan, Fatemeh Rasouli, Eppie M Yiu, Nellie Georgiou-Karistianis, Geneieve Tai, Theresa Zesiewicz, Martin B Delatycki, Louise A Corben

## Abstract

**Objective:** To identify gait and balance measures that are responsive to change during the timeline of a clinical trial in Friedreich ataxia (FRDA) we administered a battery of potential measures three times over a 12-month period.

**Methods:** Sixty-one ambulant individuals with FRDA underwent assessment of gait and balance at baseline, six months and 12 months. Outcomes included: GAITRite® spatiotemporal gait parameters; Biodex Balance System Postural Stability Test (PST) and Limits of Stability; Berg Balance Scale (BBS); Timed 25 Foot Walk Test; Dynamic Gait Index (DGI); SenseWear MF Armband step and energy activity; and the Friedreich Ataxia Rating Scale Upright Stability Subscale (FARS USS). The standardised response mean (SRM) or correlation coefficients were reported as effect size indices for comparison of internal responsiveness. Internal responsiveness was also analysed in subgroups.

**Results:** SenseWear Armband daily step count had the largest effect size of all the variables over six months (SRM=-0.615), while the PST medial-lateral index had the largest effect size (SRM=0.829) over 12 months. The FARS USS (SRM=0.824) and BBS (SRM=-0.720) were the only outcomes able to detect change over 12 months in all subgroups. The DGI was the most responsive outcome in children, detecting a mean change of −2.59 (95% CI −3.52 to −1.66, *p*<0.001, SRM=-1.429).

**Conclusions:** The FARS USS and BBS are highly responsive and can detect change in a wide range of ambulant individuals with FRDA. However, therapeutic effects in children may be best measured by the DGI.

## Introduction

Incoordination and gait ataxia are usually the first symptoms in Friedreich ataxia (FRDA), occurring between 10-15 years of age on average^1^. Mobility and balance progressively decline, with loss of ambulation approximately 10-15 years after onset^2^. Multiple pharmacological trials to evaluate potential treatments are underway with primary endpoints measuring therapeutic effectiveness typically clinical ataxia rating scales or biomarkers such as frataxin levels^3^. However, with these endpoints, power analyses indicate trial durations of over 1-2 years for an effect that slows disease progression^3^.

Given the profound impact mobility loss has on all domains of quality of life^4,5^, sensitive endpoints measuring mobility and balance are critical to ensure clinically meaningful benefit. The Timed 25 Foot Walk Test (25FWT)^6^ is the most common mobility outcome used in trials in FRDA. However, it appears less responsive than the Scale for the Assessment and Rating of Ataxia (SARA) and the Friedreich Ataxia Rating Scale (FARS)^7^ indicating it may not detect subtle change.

The aim of this study was to compare the responsiveness of gait and balance outcomes to disease progression over 12 months in ambulant individuals with FRDA. Instrumented and clinical outcomes were chosen based on their ability to detect change after therapeutic interventions^8,9^ or potential utility in measuring gait, balance or real-life ambulatory activity decline in FRDA^10-12^. We hypothesised that instrumented outcomes would be more responsive than clinical outcomes of gait and balance. The secondary aim was to establish estimated sample sizes for a parallel group, year-long trial demonstrating a 50% reduction in disease progression.

## Materials and Methods

### Study Design and Setting

This was a prospective longitudinal study, measuring gait and balance of ambulatory individuals with FRDA over 12 months. Gait and balance measurements were undertaken at the Clinical Research Centre for Movement Disorders and Gait, Kingston Centre (Australia) as well as at the University of South Florida (USF) Ataxia Research Centre (USA) and Human Functional Performance Laboratory (USA) between August 2015 and June 2018. Outcome measures were administered at baseline, six months and 12 months.

### Participants

Ambulant adults and children with FRDA were recruited through the Collaborative Clinical Research Network in FRDA (CCRN), the FRDA Global Patient Registry, the FRDA Clinic at Monash Medical Centre (Australia) and the USF Ataxia Research Centre (USA). Eligible individuals were homozygous for a GAA trinucleotide expansion in intron 1 of *FXN*^13^, able to walk independently with or without a gait aid, and aged seven years or older. Exclusion criteria included other medical conditions limiting ability to ambulate, being enrolled in a clinical trial at baseline assessment, or any anticipated change in their therapeutic regimen due to mobility decline between the baseline and six-month assessment.

### Standard Protocol Approvals, Registrations, and Patient Consents

This study was approved by the Monash Health Human Research Ethics Committee (15035A) and the USF Institutional Review Board (Pro00021414). All participants (or their parents/guardians if aged under 18 years) provided informed consent as per the Declaration of Helsinki.

### Outcome Measurements

A range of balance and mobility outcomes assessing ‘body functions’ and ‘activities and participation’ as defined by the International Classification of Functioning, Disability and Health^14^, were administered. These included clinical outcomes, laboratory-based instrumented gait and balance tests and one ‘real-life setting’ instrumented measure. Clinical rating scales were also examined to provide a reference for change.

### Clinical rating scales

Disease severity was assessed by the FARS Neurological Exam (FARS NEURO)^15^, FARS Activities of Daily Living Score (FARS ADL)^15^ and the SARA^16^. The FARS NEURO version used was scored out of 125 and includes two balance items with eyes closed^11,17^. Demographic and clinical information: age at the time of testing, age of symptom onset (age in years at which clinical symptoms of FRDA were first noticed by the individual or parents), disease duration, gait aid used and GAA1 (smaller intron 1 *FXN* GAA repeat) and GAA2 (larger intron 1 *FXN* GAA repeat) sizes, were also collected.

### Instrumented gait and balance measures

Spatiotemporal parameters of gait were measured using the GAITRite^®^ Walkway System (CIR Systems Inc., Clifton, NJ, USA). The sampling frequency was set at 100Hz. Participants were asked to walk down the 8.3m mat six times (one practice and five actual trials) each for two walking speeds: 1) self-selected preferred (preferred speed) and 2) as fast as possible (fast speed). Mean and intra-individual step variability (standard deviation) data from the five actual trials at each speed were analysed. Variables analysed were velocity, cadence, stride and step length, percentage of gait cycle in swing and in double support, heel-to-heel base of support (BOS), and stride, swing and double support time. Stride length, velocity and cadence were also normalised to height^18^.

Static and dynamic postural stability was measured by the Biodex Balance System™ SD (Biodex Medical Systems Inc., Shirley, NY) Postural Stability Test (PST) and Limits of Stability (LOS)^10^. For the PST participants were asked to balance for 30 seconds in three different conditions: 1) eyes open; 2) with visual feedback (the participant’s centre of gravity position on the platform is indicated via a cursor on a biofeedback display screen); and 3) eyes closed. A medial-lateral, anterior-posterior and overall stability index was calculated for each condition. For the LOS, participants were required to shift their weight to move their centre of gravity from a central position to eight peripheral targets and return to the central position in random order. Time taken (LOS duration) to reach all targets and the overall directional score (LOS score) were recorded. For all tests and conditions, two practice trials were conducted^19^ and a maximum of five trials were allowed to achieve three successful trials (maintaining balance during a completed trial) in each condition.

Daily walking and activity levels were measured by the SenseWear MF Armband (SenseWear) (BodyMedia Inc., Pittsburgh, PA, USA), a wireless activity monitor using a triaxial accelerometer worn over the triceps^20^. Variables measured were total energy expenditure (kcal/min), active energy expenditure (kcal/min), Metabolic Equivalent of Task (MET) levels, daily step count, daily distance walked (km), physical activity duration (min), lying down time (min) and time spent in sedentary, light, moderate, vigorous and very vigorous activity (min). Participants were required to wear the SenseWear accelerometer in their usual environment for 24 hours per day (removing for bathing only) for five consecutive typical daily living days. SenseWear data were included in the analysis if the SenseWear was worn for a ‘valid day’ (>21.6 hours/90%).

### Clinical gait and balance measures

Ambulation was measured using the 25FWT and the Dynamic Gait Index (DGI)^21^. Balance was assessed using the Berg Balance Scale (BBS)^22^ and the FARS Upright Stability Subscale (FARS USS) scored out of 36^11,17^. All clinical tests were performed according to established protocols^11,22-24^.

### Procedure

Gait and balance assessments were administered in one day. Assessments occurred in the same order to standardise the effects of fatigue: 1) GAITRite® preferred and fast walks, 2) PST and LOS, 3) BBS, 4) DGI and 5) 25FWT. The FARS NEURO (including the FARS USS), FARS ADL and SARA were administered within one week, and the SenseWear worn within 40 days of the gait and balance assessment.

### Data Analysis

Normal distribution was analysed using histograms and the Shapiro-Wilk test. The reciprocal of the 25FWT (25FWT^-1^) was calculated to account for the skewness of data^25^. Paired sample t-tests (or Wilcoxon sign rank tests in the event of non-parametric data and sample sizes n<30) with 95% confidence intervals were utilised to determine significant change between: 1) baseline and six-month visit and 2) baseline and 12-month visit. The standardised response mean (SRM) as mean score change/SD of score change was reported as the effect size index to enable a comparison of internal responsiveness between the scales^26^. Values of 0.20, 0.50 and 0.80 or greater were the criterion for small, moderate, and large changes respectively^27^. Correlation coefficients (r=z/√ n where n = total number of observations) were used as the effect size index for non-parametric data^28^.

Subgroup analyses of internal responsiveness were conducted using the same method to ascertain differences between: a) children and adults; b) those ambulant with and without an aid; and c) those with typical onset (onset of symptoms <25 years of age) and Late Onset Friedreich Ataxia (LOFA) (onset of symptoms ≥ 25 years of age^29^). To establish estimated sample sizes to demonstrate a reduction of 50% in disease progression over a 12-month period, sample size calculations were conducted using a 2-tailed type I error of < 5% with a power of 80%. Significance was recorded as *p*<0.05. The analysis was performed using STATA (StataCorp. 2015. Stata Statistical Software: Release 15. College Station, TX: StataCorp LP).

### Data Availability Statement

Data not published within this article will be shared after approval by the Friedreich’s Ataxia Research Alliance, USA, and Ethics Review Boards.

## Results

### Participants

Sixty-one participants met the study criteria, were enrolled and completed the baseline assessment. Participant demographics and clinical data are summarised in Table 1. There were 43 adults and 18 children; 38 individuals with typical onset FRDA and 23 with LOFA; and 50 individuals ambulant without an aid and 11 ambulant with an aid at baseline. There were no significant differences in demographics or clinical parameters across the two sites (data not shown). Participants requiring a gait aid during the gait assessments had a disease duration of 16.44 ± 5.65 (mean ± SD) years, while those not requiring an aid had a disease duration of 9.93 ± 6.70 years, *p*=0.004. Fifty-eight (95.1%) participants completed the six-month visit 185.4 ± 18.7 (range, 151-230) days after baseline. Fifty-four (88.5%) participants returned for the 12-month visit, 372.5 ± 24.5 (287-422) days after baseline. Reasons for withdrawal included: medical restrictions unrelated to FRDA (n=4); school schedule (n=1); pregnancy (n=1); and declined (n=1).

**Table 1.** Baseline characteristics of participants.

| Clinical Parameters | Whole cohort (n=61) |  | Children (n=18) |  | Adults (n=43) |  |
| --- | --- | --- | --- | --- | --- | --- |
|  | Mean (SD) or Ratio | Range | Mean (SD) or Ratio | Range | Mean (SD) or Ratio | Range |
| **Gender (male/female)** | 36/25 |  | 12/6 |  | 24/19 |  |
| **Age at assessment (years)** | 28.09 (13.48) | 7.01-54.59 | 13.15 (2.95) | 7.01-17.65 | 34.34 (10.97) | 18.14-54.59 |
| **Age of onset (years)** | 16.98 (9.90) | 2-42 | 7.11 (3.74) | 2-16 | 21.12 (8.66) | 5-42 |
| **Type of onset (typical/late)** | 38/23 |  | 18/0 |  | 20/23 |  |
| **GAA1** | 555.02 (256.42) | 55-1050 | 806.11 (135.33) | 514-1050 | 449.91 (219.52) | 55-1050 |
| **GAA2** | 875.67 (259.29) | 182-1555 | 961.00 (196.87) | 750-1555 | 839.95 (275.55) | 182-1266 |
| **Disease duration (years)** | 11.11 (6.96) | 0.66-36.08 | 6.03 (2.36) | 0.66-10.70 | 13.23 (7.16) | 3.15-36.08 |
| **FARS NEURO** | 47.47 (13.42) | 12.0-75.0 | 48.62 (14.73) | 12.0-75.0 | 47.02 (13.03) | 15.5-73.5 |
| **SARA** | 12.17 (4.79) | 2.0-24.5 | 12.22 (5.50) | 2.0-24.5 | 12.15 (4.53) | 4.5-21.0 |
| **Gait aid (without/with)** | 50/11 |  | 16/2 |  | 34/9 |  |

Abbreviations: GAA1: GAA repeat size of the smaller FXN allele; GAA2 = GAA repeat size of the larger FXN allele; FARS NEURO: Friedreich Ataxia Rating Scale Neurological Exam; SARA: Scale for the Assessment and Rating of Ataxia.

Fifty out of 61 (82.0%), 47/58 (81.0%) and 41/54 (75.9%) participants were able to complete the GAITRite® assessment without a gait aid at the baseline, six-month and 12-month assessment, respectively. Two participants were unable to walk without physical assistance at the 12-month visit and did not complete the GAITRite® assessment. At baseline, 58/61 (95.1%) participants completed the Biodex PST eyes open and visual feedback conditions and 28/61 (45.9%) completed the eyes closed condition. At 12 months this number decreased to 49/54 (90.7%) completing the visual feedback condition, 47/54 (87.0%) the eyes open condition and 18/54 (33.3%) the eyes closed condition. SenseWear data was collected from 32, 25 and 23 participants at baseline, six months and 12 months respectively. Missing SenseWear data were primarily due to participants’ late return of the SenseWears and a subsequent delay in distribution to other participants.

### Clinical outcomes

Histograms of clinical outcomes at baseline are presented in Figure 1. At six months, the BBS, DGI and FARS USS all detected decline with small effect sizes; however, there was no significant change in the 25FWT^-1^, FARS NEURO, FARS ADL or SARA. Over 12-months, the FARS USS was the most responsive clinical outcome (SRM=0.824). See Table 2 for details of clinical outcomes.

**Table 2.** Change in clinical gait and balance measures.

| *Variable* | *Baseline*<br>*Mean (SD)* | *Six Month*<br>*Mean (SD)* | *Six Month*<br>*Change*<br>*Mean [95% CI]* | *t-test<sup>a</sup>, SRM* | *12 Month*<br>*Mean (SD)* | *12 Month Change*<br>*Mean [95% CI]* | *t-test<sup>a</sup>, SRM* |
| --- | --- | --- | --- | --- | --- | --- | --- |
| BBS | 42.95<br>(11.78) | 40.5 (12.63) | -1.91 [-2.97, -0.86] | $t(57)=-3.635$, $p=0.001$, SRM=-0.477 | 39.54<br>(13.37) | -2.94 [-4.96, -1.83] | $t(53)=-5.290$, $p<0.001$, SRM=-0.720 |
| DGI | 13.75 (5.73) | 12.84 (5.74) | -0.67 [-1.15, -0.19] | $t(57)=-2.799$, $p=0.007$, SRM=-0.368 | 12.26 (5.46) | -1.54 [-2.14, -0.93] | $t(53)=-5.086$, $p<0.001$, SRM=-0.692 |
| 25FWT<sup>-1</sup> | 0.14 (0.06) | 0.14 (0.07) | 0.01 [-0.00, 0.01] | $t(54)=1.421$, $p=0.161$, SRM=0.192 | 0.13 (0.07) | -0.01 [-0.02, 0.01] | $t(50)=-1.016$, $p=0.315$, SRM=-0.142 |
| FARS USS | 19.22 (6.07) | 20.50 (5.93) | 1.09 [0.46, 1.71] | $t(56)=3.469$, $p=0.001$, SRM=0.459 | 21.19 (6.04) | 1.85 [1.23, 2.47] | $t(52)=5.996$, $p<0.001$, SRM=0.824 |
| SARA | 12.17 (4.79) | 13.07 (4.40) | 0.66 [-0.14, 1.45] | $t(57)=1.646$, $p=0.105$, SRM=0.216 | 13.32 (4.51) | 1.03 [0.29, 1.77] | $t(52)=2.800$, $p=0.007$, SRM=0.385 |
| FARS NEURO | 47.47<br>(13.42) | 49.43<br>(12.56) | 1.35 [-0.78, 3.48] | $t(56)=1.269$, $p=0.210$, SRM=0.168 | 49.07<br>(12.77) | 1.51 [0.05, 2.98] | $t(52)=2.071$, $p=0.043$, SRM=0.284 |
| FARS ADL | 11.08 (4.60) | 12.27 (4.46) | 1.01 [-0.18, 2.21] | $t(34)=1.728$, $p=0.093$, SRM=0.292a | 12.82 (5.02) | 1.77 [0.72, 2.82] | $t(30)=3.450$, $p=0.002$, SRM=0.620 |

<sup>a</sup>Paired sample t-tests were used for the analysis. Abbreviations: 25FWT<sup>-1</sup>: reciprocal of the Timed 25 Foot Walk Test; BBS: Berg Balance Scale; CI: Confidence Interval; DGI: Dynamic Gait Index; FARS ADL: Friedreich Ataxia Rating Scale Activity of Daily Living score; FARS NEURO: Friedreich Ataxia Rating Scale Neurological Exam; FARS USS: Friedreich Ataxia Rating Scale Upright Stability subscale; SARA: Scale of the Assessment and Rating of Ataxia; SD: standard deviation; SRM: standardised response mean.

**Figure 1.**
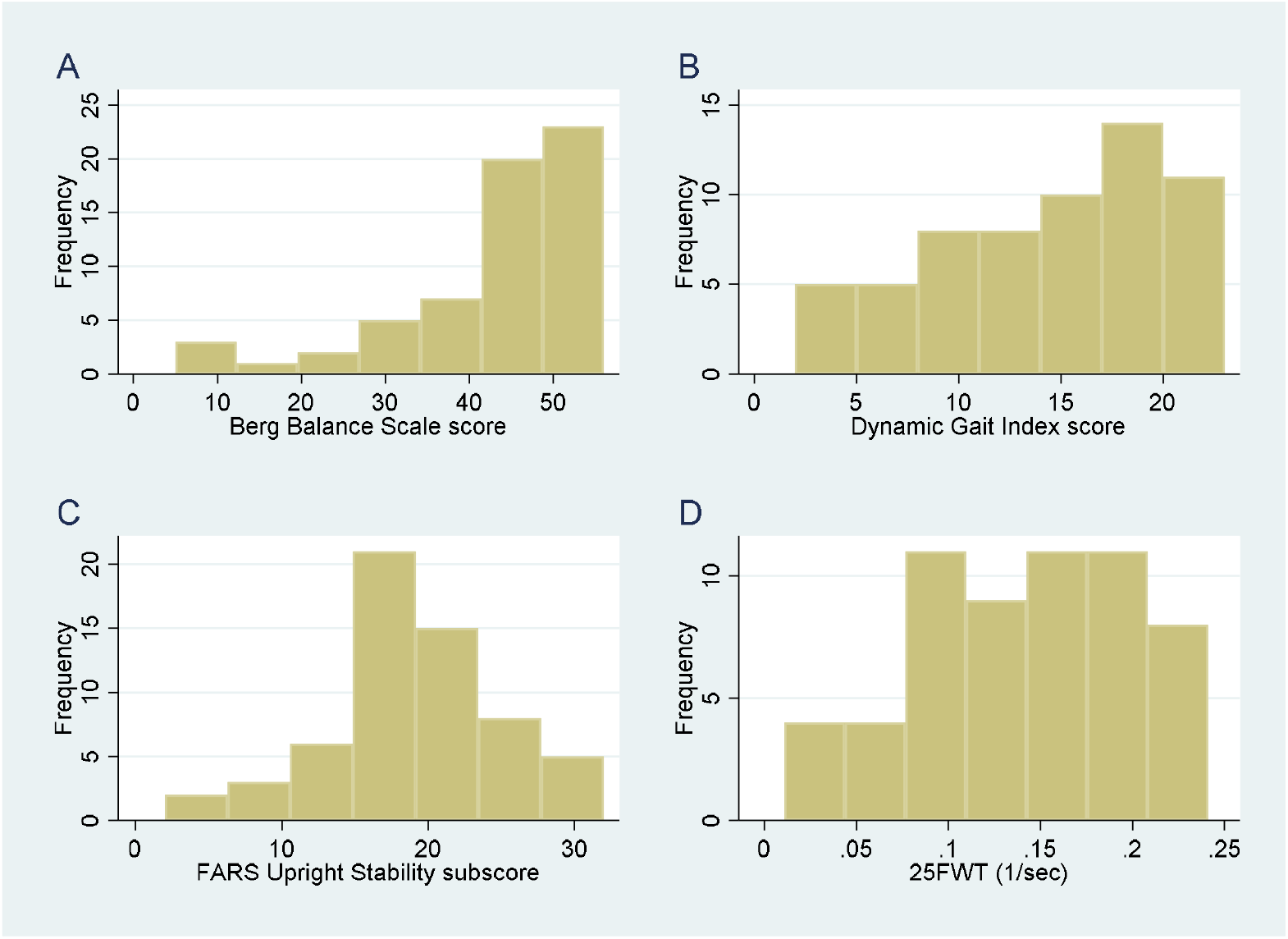
Histograms of clinical gait and balance outcomes. Figure Legend: Histograms of the A. Berg Balance Scale; B. Dynamic Gait Index; C: Friedreich Ataxia Rating Scale Upright Stability Subscale; D. Reciprocal of the Timed 25 Foot Walk Test. Abbreviations: 25FWT: Timed 25 Foot Walk Test; FARS: Friedreich Ataxia Rating Scale.

### GAITRite® spatiotemporal parameters

Participants took a mean of 61.69 ± 19.48 and 53.99 ± 15.56 steps on the GAITRite® walkway during preferred and fast speed, respectively. There was a significant difference in velocity between the preferred and fast speed condition at each visit. At baseline, only preferred speed BOS, preferred speed BOS variability and fast speed BOS variability were normally distributed. Stride length during fast speed was the only spatiotemporal gait parameter responsive to change over six months, reducing by a mean 2.82 cm (95% CI −5.43 to −0.21; *p*=0.035, SRM=-0.298). Over 12 months, fast speed velocity and cadence had the greatest effect sizes of the spatiotemporal gait parameters: velocity decreased by a mean 10.55 cm/s (95% CI - 15.27 to −5.82; *p*<0.001, SRM=-0.641) and cadence decreased by 5.30 steps/min (95% CI −7.73 to −2.87; *p*<0.001, SRM=-0.626). See Figure 2 for change in mean spatiotemporal gait parameters. The only parameter of variability to show significant change over 12 months was preferred speed BOS variability (0.40cm [mean change], 95% CI 0.05 to 0.75; *p*=0.024, SRM=0.322).

**Figure 2.**
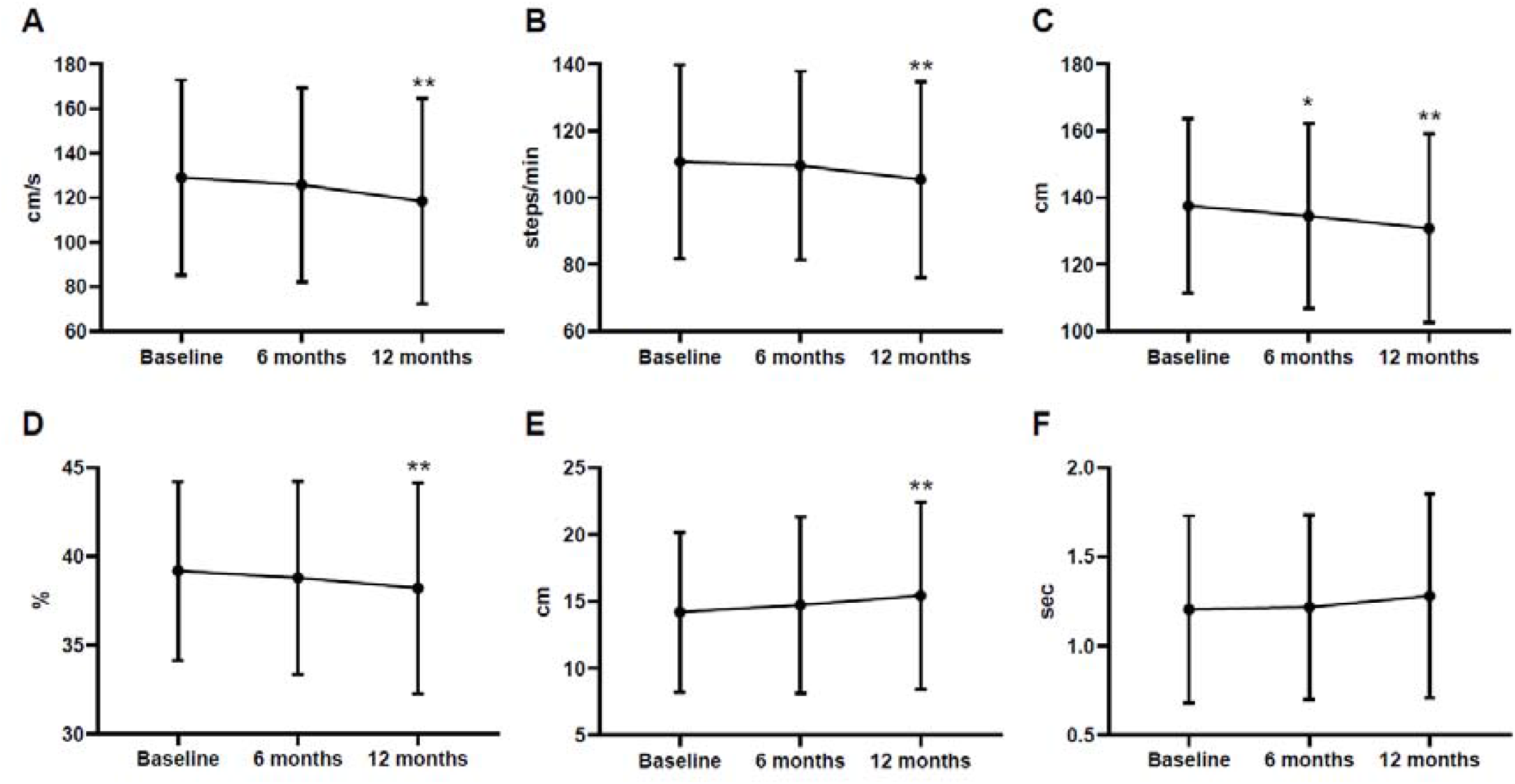
Mean (SD) in fast speed mean spatiotemporal gait parameters during each visit. Figure Legend: Mean (SD) in A. Velocity; B. Cadence; C. Stride length; D. Swing as a percentage of the gait cycle; E. Heel-to-heel base of support; and F. Stride time, during the fast speed condition at each visit. Data presented is not normalised for height. Significant change over six months is indicated by one asterisk (*p*<0.05). Significant change over 12 months is indicated by two asterisks (*p*<0.05).

### Biodex Balance System variables

At baseline, only PST anterior-posterior and overall indices with eyes closed and LOS score were normally distributed. Over six months the sole PST index to identify a decline was the anterior-posterior index with eyes closed, scores increased by 0.36 (95% CI 0.04 to 0.68; *p*=0.030, SRM=0.510). Over six months, the LOS score increased by 2.40 (95% CI: 0.61 to 4.18; *p*=0.010) and the LOS duration decreased by 6.47 seconds (95% CI −10.94 to −1.99; *p*=0.006) suggesting an improvement in dynamic stability.

Over 12 months, the PST medial-lateral index with eyes closed increased by 0.33 (95% CI 0.13 to 0.54, *p*=0.004, SRM=0.829), indicating decline in postural stability. The PST overall index with eyes closed also significantly increased by 0.37 (95% CI 0.02 to 0.72; *p*=0.041, SRM=0.540). The six-month change detected by the PST anterior-posterior index with eyes closed was not present at 12 months. There was no significant change in the eyes open or visual feedback conditions. The LOS variables continued to indicate an improvement in stability at 12 months: LOS score increased by 2.42 (95% CI 0.06 to 4.78; *p*=0.045) and LOS duration decreased by 7.33 seconds (95% CI −13.23 to −1.44; *p*=0.016).

### SenseWear Armband accelerometer variables

Participants wore the SenseWear 97.60 ± 0.01% per day, for 4.38 ± 0.96 valid days. There were several outliers present in the SenseWear data (Figure 3). Over six months, physical activity duration reduced by 30.22 min (95% CI −56.86 to −3.59; *p*=0.028, SRM=-0.424); distance walked reduced by 0.96 km (95% CI −1.67 to −0.25; *p*=0.010, SRM=-0.506) and daily step count decreased by 1037.64 steps (95% CI - 1667.48 to −407.80; *p*=0.002, SRM=-0.615). Over 12 months, the significant decline continued only in daily step count (baseline [median] 3898.00 vs 12-months: 2849.20 steps; z=-2.19; *p*=0.029, r=-0.297). When the analysis was repeated with the outliers removed, there was also a significant reduction in distance walked over 12 months (baseline: 3.19 vs 12-months: 1.56 kms, z=-2.121, *p*=0.034, r=0.294).

**Figure 3.**
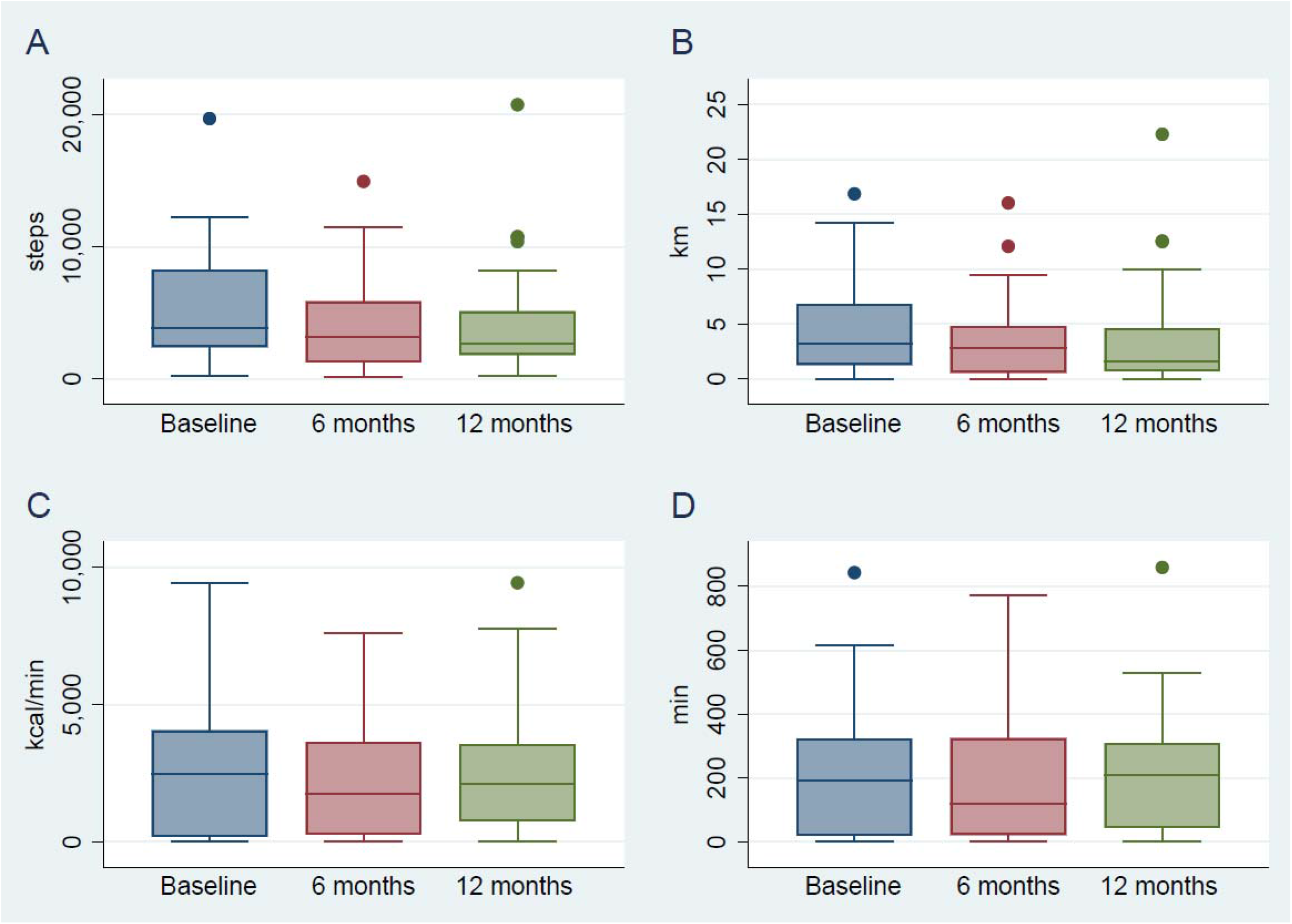
Box and whisker plots of SenseWear Armband accelerometer data. Figure Legend: Box and whisker plots of the SenseWear accelerometer at each visit for participants completing all three assessments (n=26). A. Daily Step Count; B. Daily Distance Walked; C. Active Energy Expenditure; D. Physical Activity Duration. Boxes represent the median value (middle line) and inter-quartile range (25%-75%). Whiskers extend to the lower and upper adjacent values. Dots represent outliers.

### Children compared to adults

In children (n=18), the DGI was the most responsive measure, detecting a mean change of −2.59 (95% CI −3.52 to −1.66, p<0.001, SRM=-1.429). The BBS (SRM=-0.949) and FARS USS (SRM=1.055) also had large effect sizes over 12 months in children. Comparatively in adults (n=43), the DGI decreased by 1.05 (95% CI −1.80 to −0.30; *p*=0.007, SRM=-0.469). The most responsive outcomes for adults were the FARS USS (SRM=0.725) and PST medial-lateral index with eyes closed (SRM=0.743). The SARA (95% CI 0.10 to 1.93; *p*=0.031, SRM=0.374) and FARS ADL score (95% CI 0.56 to 2.94; *p*=0.006, SRM=0.652) detected a change in adults, but not in children, over 12 months.

Several mean spatiotemporal gait parameters during fast speed demonstrated significant changes in both adults and children (Figure 4). However, BOS variability was the only preferred speed parameter to detect change in children (z=2.533, *p*=0.011, r=0.434). Fast speed stride length variability demonstrated a decrease of 1.12cm (95% CI −2.18 to −0.06; *p*=0.040, SMR=-0.367) in adults and an increase of 2.00cm (95% CI 0.57 to 3.43; *p*=0.010, SMR=0.775) in children. No other variable demonstrated a different pattern of change in children compared to adults.

**Figure 4.**
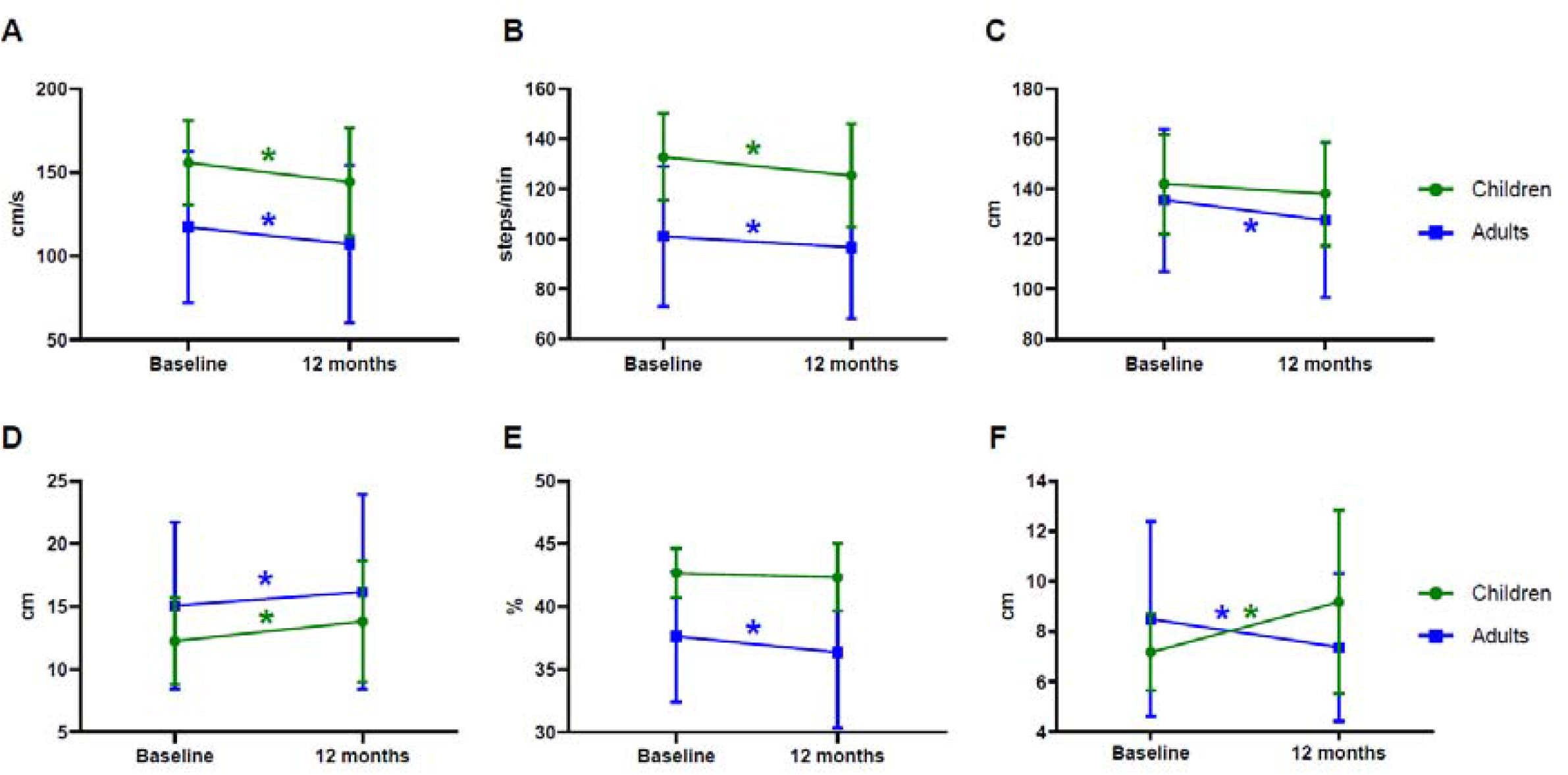
Twelve-month change in fast speed spatiotemporal gait parameters in adults and children. Figure Legend: Mean (SD) in A. Velocity; B. Cadence; C. Stride length; D. Heel-to-heel base of support; E. Swing as a percentage of the gait cycle; F. Stride length variability in the fast speed condition at the baseline and 12-month visits. Data presented is not normalised for height. Significant change over 12 months is indicated by the asterix.

### People ambulant with and without an aid

Only the BBS, FARS USS and preferred speed BOS variability detected a significant change in both groups at 12 months. For individuals ambulating without an aid (n=50), the PST medial-lateral index with eyes closed (SRM=0.829) had the largest effect size, followed by the DGI (SRM=-0.812). In this subgroup, all mean fast speed parameters, unadjusted for height, indicated a significant decline. However, BOS variability was the only preferred speed parameter to detect a change, increasing by 0.43cm (95% CI 0.04 to 0.83; *p*=0.033, SMR=0.344).

For individuals ambulating with an aid (n=11), preferred speed BOS variability (SRM=0.831), BBS (SRM=-0.913), FARS USS (SRM=0.937), PST medial-lateral index with visual feedback (SRM=1.029) and time spent in sedentary activity (SRM=-1.616) had large effect sizes. The FARS ADL significantly increased by 1.87 (95% CI 0.70 to 3.03; *p*=0.003, SRM=0.694) and daily step count decreased (z=-2.312, *p*=0.021, SRM=-0.341) in individuals ambulating without an aid but not in those using an aid, while the 25FWT^-1^ significantly decreased by −0.01 (95% CI −0.03 to 0.00; *p*=0.042, SRM=-0.704) only for those using an aid.

### Typical compared to Late Onset Friedreich ataxia

For individuals with typical onset FRDA (n=38), the FARS USS (SRM=0.983), DGI (SRM=-0.900), cadence (SRM=-0.976) and velocity (SRM=-0.872) at fast speed, and FARS ADL (SRM=0.803) had large effect sizes for change over 12 months. The PST medial-lateral index with eyes closed (SRM=1.377) was the most responsive measure over 12 months in individuals with LOFA (n=23) but did not detect a change in individuals with typical onset. Outcomes detecting a change in both subgroups were the BBS (LOFA [mean change]: −2.45; 95% CI −4.36 to −0.54; *p*=0.014, SRM=-0.602; typical onset: −3.24, 95% CI −4.68 to −1.79; *p*<0.001, SRM=-0.783) and the FARS USS (LOFA: 1.35, 95% CI 0.27 to 2.43; *p*=0.017, SRM=0.586; typical onset: 2.16, 95% CI 1.38 to 2.94; *p*<0.001, SRM=0.983). The FARS NEURO increased by 2.64 (95% CI 0.62 to 4.66, *p*=0.012, SRM=0.464) and the SARA increased by 1.31 (95% CI 0.38 to 2.24; *p*=0.007, SRM=0.491) in individuals with typical onset FRDA but not in individuals with LOFA.

### Sample size estimations

The estimated sample sizes required to achieve a reduction of 50% in disease progression over a 12-month period for the SARA and FARS NEURO were 432 and 794 participants per group, respectively. In comparison, the FARS USS produced an estimated sample size of 95 participants per group. See Table 3 for estimated sample sizes for variables with large effect sizes.

**Table 3.** Estimated sample sizes for a one-year trial.

| Variable | Cohort<sup>a</sup> | Estimated Sample Size<br>(participants per group) |
| --- | --- | --- |
| SenseWear Time spent in Sedentary Activity | Ambulating with gait aid | N/A<sup>b</sup> |
| Dynamic Gait Index | Children | 32 |
| PST medial-lateral index with eyes closed | LOFA | 36 |
| Mean cadence at fast speed | Children | 51 |
| FARS Upright Stability subscale | Children | 58 |
| PST medial-lateral index with visual feedback | Ambulating with gait aid | 60 |
| FARS Upright Stability subscale | Typical onset | 67 |
| Mean cadence at fast speed | Typical onset | 68 |
| Berg Balance Scale | Children | 71 |
| Mean velocity at fast speed | Children | 72 |
| FARS Upright Stability subscale | Ambulating with gait aid | 73 |
| Berg Balance Scale | Ambulating with gait aid | 77 |
| Dynamic Gait Index | Typical onset | 79 |
| Mean velocity at fast speed | Typical onset | 84 |
| FARS Upright Stability subscale | Whole cohort | 95 |
| BOS variability at preferred speed | Ambulating with gait aid | 96 |
| Dynamic Gait Index | Ambulating without an aid | 98 |
| Mean stride time at fast speed | Child | 98 |
| FARS Activities of Daily Living score | Typical onset | 99 |
| PST medial-lateral index with eyes closed | Ambulating without an aid | 100 |
| PST medial-lateral index with eyes closed | Whole cohort | 100 |
| SARA<sup>c</sup> | Whole cohort | 432 |
| FARS NEURO<sup>c</sup> | Whole cohort | 794 |

<sup>a</sup>There were no variables with large effect sizes in adults. <sup>b</sup>Time in sedentary activity decreased over 12 months in individuals therefore the estimated sample size is not applicable. <sup>c</sup>The SARA and FARS NEURO are displayed for comparative purposes and did not have large effect sizes. Abbreviations: BOS: heel-to-heel base of support; FARS: Friedreich Ataxia Rating Scale; FARS NEURO: Friedreich Ataxia Rating Scale Neurological Exam; LOFA: late onset Friedreich ataxia; medial-lateral: medio-lateral; PST: Postural Stability Test; SARA: Scale for the Assessment and Rating of Ataxia.

## Discussion

This study compared the responsiveness of instrumented and clinical gait and balance outcomes over six and 12 months in ambulant individuals with FRDA. Variables from the instrumented devices, SenseWear and Biodex Balance System™ SD were more responsive to disease trajectory than clinical outcomes, with SenseWear daily step count and the PST medial-lateral index with eyes closed the most responsive outcomes over six months and 12 months, respectively. However, utilisation of these outcomes for clinical trials may be limited due to large variability (SenseWear daily step count) and significant floor effects (66.6% of participants were unable to complete the PST medial-lateral index with eyes closed at 12 months). In contrast, the FARS USS had a large effect size for change over 12 months, demonstrated consistent change in children, adults and in individuals with typical and late onset FRDA, and did not exhibit significant floor or ceiling effects.

Balance outcomes were more responsive to 6 and 12 months of natural disease progression than spatiotemporal gait parameters and accelerometry outcomes in this study. In particular, the significant decline identified by the PST eyes closed condition, though not completed by more half the participants, highlighted an increase in visual reliance for static stability early in the disease. The BBS and FARS USS include items that measure balance with both eyes open and eyes closed, which may have contributed to their sensitivity. Both also include items of dynamic and static stability, further expanding the breadth of their evaluation in ambulant individuals. This was reflected in their ability to detect change in all subgroups in this study. Conversely, it may be that certain items in these scales (such as single leg or tandem stance) produced large changes in this cohort due to participants only scoring maximum or minimum values (reflecting either ability or inability to complete the task) suggested by the binomial distributions previously seen in six of the FARS USS items^30^. The DGI, measuring dynamic stability during walking, a slightly different construct to the FARS USS and BBS, was also responsive in four of the six subgroups; however, three items exhibited floor effects when participants were dependent on a gait aid to ambulate, potentially reducing its overall responsiveness in this cohort.

The potential for the SenseWear to be sensitive to change may have been limited by high levels of variability between and within subjects’ performance. Over a 12-month period, participant-related factors, such as seasonal influences on activity, changes in lifestyle or residence, enjoyment of physical activity, motivation and adherence to study protocols, may have influenced activity levels. Technology-related factors, such as triceps placement of the SenseWear accelerometer, detection of lower ambulation speeds^31^, gait aid limitations^32^ and ability to download data remotely may also have impacted data collection accuracy. Furthermore, not all participants completed this evaluation, resulting in a smaller sample size for the SenseWear variables with an increased risk of a false-negative findings^33^. Measurement of real-life activity is important to understand the true impact of therapeutic interventions. Promising studies have suggested use of body-worn sensors that capture variables not influenced by ‘condition-dependent’ or external factors, improve the accuracy and responsiveness of ambulatory measures while reducing the burden on participants^34,35^. Moreover, establishing the best measurement duration, balancing burden, validity and reliability, is important^35^. The responsiveness of the SenseWear seen in this study, even with the potential limitations, indicates merit in further evaluation of real-life ambulation. Opportunities to manage the influencing factors will ensure clinically and ecologically^34^ meaningful evaluation of ambulation.

An unexpected finding in this study was detection of an improvement in the LOS score and duration. This contrasted with the decline detected by the PST eyes closed condition and clinical balance measures. The LOS measures dynamic stability, which has different system demands to those required for static postural control and previous studies have shown a poor relationship between static and dynamic balance tests^36^. It is unlikely that cerebellar control of dynamic stability would improve given the neurodegenerative nature of FRDA^1^; however, individuals with FRDA may be able to increase cognitive regulation of their performance during the LOS to maintain upright stability, until their attentional capacity has been reached^37^. This apparent improvement in dynamic stability is consistent with the reduction in stride length variability that was seen over time in adult participants. Conversely, LOS changes may be due to a learning effect. Two practice trials were performed in this study as standard practice to maximise test-retest reliability^19^. However, for individuals with FRDA fatigue levels may have altered this mitigating process. This highlights the requirement for reliability and measurement error testing specifically in FRDA. Moreover, including both dynamic and static stability tests appears critical to ensure balance and stability are measured accurately.

Although the clinical and instrumented balance outcomes were more responsive than the outcomes specifically measuring gait and accelerometry outcomes, several changes in spatiotemporal parameters were identified over six and 12 months. Preferred speed BOS variability was responsive for children, individuals with typical onset, and those ambulating without and with a gait aid over 12 months. BOS variability is thought to be an indicator of lateral balance control^38^ and is dependent on visual-vestibular feedback^39^. As an adaptation to gain stability during walking, too much or too little BOS variability can be an early predictor of falls^40^. Moreover, BOS variability appears not to be altered by gait aid use^41^ nor is dependent on gait speed^42^, resulting in its utility as an outcome measure for individuals with a wide spectrum of walking ability. The second pattern of changes was similar to those seen in the pilot study by Zesiewicz and colleagues^11^, where velocity, cadence and stride length, when walking at maximum speed, showed decline over 12 months. As cadence, stride length and velocity are interrelated^43^, decline may primarily be a consequence of reduced walking speeds to adapt for loss of balance or sensory impairment^44,45^. Lastly, in contrast to the findings of Zesiewicz and colleagues^11^, stride length variability did not significantly increase in the whole cohort. This is surprising as gait variability is a significant feature of FRDA^10^ and has been shown to increase when speeds decrease^42^. Analysis of subgroups found stride length variability significantly increased in children and significantly decreased in adults. This unexpected finding in adults may have been reflected in previous cross-sectional studies identifying an inconsistent relationship between stride length variability and disease duration^10,12^ However, the clinical significance of this is uncertain. In contrast, the increase in stride length variability in children has significant clinical relevance given gait should be maturing with decreasing variability until adulthood^46^. Considering these gait changes, to provide a comprehensive evaluation of spatiotemporal changes in individuals with FRDA, an assessment of preferred speed BOS variability, and maximum safe speed velocity and stride length variability should be undertaken. Moreover, further work to develop a compound measure, similar to that utilised by Ilg and colleagues^34^, may provide an even more sensitive measure of gait deterioration.

Several limitations should be noted with the interpretation of the study findings. First, test-retest reliability of the balance outcomes has not been established in individuals with FRDA. Many individuals with FRDA comment on their fluctuating physical ability related to time of day, fatigue and current activity levels. However, perceived fluctuation may not be present in laboratory-based gait testing with strong test-retest reliability of spatiotemporal gait parameters seen in ambulatory individuals with FRDA over a 48-hour period^47^. In addition, the BBS and DGI are both subjective balance outcomes and have not had their psychometric properties evaluated in individuals with FRDA. Although these clinical measures have shown good reliability in other neurodegenerative conditions^48^ and were responsive to coordination training in individuals with degenerative ataxia^8,9^, the clinical features associated with FRDA may not facilitate an easy interpretation for scoring. Further clarification of item descriptions and a clear definition of ‘supervision’ may further increase the responsiveness of these outcomes. The second limitation of this study relates to the outcomes analysed. Straight walking tasks and balance outcomes may not have enough complexity to challenge the dentate nucleus, a significant site of cerebellar pathology in FRDA^2^. Dual tasking, obstacle clearance and multi-directional walking may detect neurodegeneration in FRDA more precisely by assessing neurocognitive, visual involvement and higher level processing in gait^49^. Nevertheless, the decline seen in the PST eyes closed condition and spatiotemporal parameters suggests sensory and proprioceptive impairment^45,50^ may be the primary neuropathology affecting postural stability in individuals with FRDA.

Utilising responsive outcomes reduces the burden of study participation by decreasing the number of participants required and reducing the duration of participation. This study has provided evidence for the use of clinical balance outcomes, the FARS USS and BBS, in trials involving ambulant individuals with FRDA. Future work is required to establish clinically meaningful change in these outcomes, establish test-retest reliability and confirm if changes are consistent across longer timeframes to ensure appropriate follow up in trials. Although there were limitations with the instrumented outcomes, their potential as highly responsive outcomes in this cohort is apparent. Addressing their limitations and ensuring they can measure change in individuals across the spectrum of gait and balance ability will improve their utility and responsiveness. This will not only reduce the sample size required but will increase the recruitment pool, improving generalisability and feasibility of clinical and therapeutic trials for individuals with FRDA.

## Acknowledgements

We wish to thank all the participants who have given their time for this project. The authors would also like to thank the Kingston Centre Physiotherapy Department, Monash Health, Melbourne, and Clinical Research Centre for Movement Disorders and Gait, Monash Health, Melbourne, for their support during the study. This study is funded by the Friedreich’s Ataxia Research Alliance (USA) (FARA), PTC Therapeutics and Voyager Therapeutics, as a part of FARA’s Biomarker Consortium.

